# A 3D framework for delimiting a polyploid complex in *Rorippa* (Brassicaceae): combining trait evolution, herbarium records, and machine learning

**DOI:** 10.1101/2025.06.26.661803

**Authors:** Ting-Shen Han, Jun-Xian Lv, Yao-Wu Xing

## Abstract

Species delimitation in polyploid complexes remains a fundamental challenge due to pervasive morphological overlap and genomic redundancy. We examined the *Rorippa dubia–indica* complex (Brassicaceae), a polyploid lineage comprising tetraploid and hexaploid taxa. We developed an integrative 3D framework (Delimitation, Distribution, and Decoding) that synthesizes spatiotemporal data from field (2017–2020; *n* = 3,136) and herbarium (1893–2021; *n* = 2,015) collections to diagnose misidentification, model distributions, and reconstruct classification criteria used in polyploid complexes. Morphological traits with varying degrees of plasticity were evaluated under controlled conditions to identify stable diagnostic characters. Seed arrangement, petal number, and genome size or ploidy level exhibited clear interspecific differentiation. Phylogenomic analyses based on chloroplast genomes further defined species boundaries clarified by these taxonomic traits. We then revised herbarium specimens and applied machine learning classification models to assess the extent of specimen misidentification and to recover the trait-based rationale behind species assignments. Initial misidentification rates reached 12–50% across virtual or physical specimens, largely due to reliance on plastic traits. These errors substantially distorted spatial distribution models and future climate projections. Our findings underscore the need for secondary specimen evaluation and demonstrate the importance of integrating morphologic and phylogenetic inference with machine learning tools to resolve morphologically overlapping polyploid complexes. This approach offers direct applications for biodiversity assessment, evolutionary research, and conservation planning.

## Introduction

Accurate species delimitation is fundamental to taxonomy, evolutionary biology, and conservation science (de Queiroz 2007). Traditionally, species boundaries have been inferred from readily observable morphological traits. However, trait plasticity and convergence can obscure true species boundaries, especially in lineages shaped by polyploidization (i.e., whole-genome duplication) or hybridization (Hörandl 2022; Chambers et al. 2025). In particular, polyploid complexes represent a continuum of evolutionary units connecting polyploids with their progenitors (Stebbins 1971), often characterized by overlapping morphological and eco-genetic features. These overlaps can result from shared ancestry, genomic redundancy, recurrent formation, or cytotype interaction (e.g., inter-ploidy introgression), making the delimitation of species within such complexes especially challenging (Soltis et al. 2010; Blischak et al. 2018). Taxonomic uncertainties may further distort species distribution models, misinform conservation planning or public concerns, and compromise evolutionary analyses (Wiens 2007; Hong 2016; Lughadha et al. 2018). Therefore, a critical step in delimiting taxonomically complex taxa is the secondary reassessment of population-level specimens from herbarium or field collections, using stable diagnostic traits, robust phylogeny, and innovative methods (Edwards & Knowles 2014; Hong 2025).

Recently, machine learning tools have been increasingly applied to improve species delimitation (Karbstein et al. 2024; Salles & Domingos 2025). Decision tree–based classifiers, such as conditional inference trees or classification and regression trees, are particularly useful in binary delimitation tasks. These methods identify diagnostic thresholds among multivariate traits and reveal hidden patterns that underlie taxonomists’ appraisal (Smith & Carstens 2020). By evaluating trait importance and classification rationale, machine learning models can reconstruct and audit past taxonomic decisions, thereby expose the limitations of traditional identification criteria (Edwards & Knowles 2014). In polyploid complexes, where subtle morphological differences are compounded by robust plasticity and reticulate genealogy, such tools may offer complementary options to refine taxonomic boundaries.

To achieve these objects, we introduce a 3D framework—Delimitation, Distribution, and Decoding—that unifies laboratory trait assessment, herbarium specimen review, and machine learning–based classification across spatiotemporal scales (Fig. 1). This framework enables cross-validation of species boundaries and provides a transparent pipeline for diagnosing misidentification, modeling ecological distributions, and reconstructing classification criteria used in morphologically overlapping species complexes. It incorporates fine-scale phenotyping from contemporary populations grown under controlled conditions, large-scale review of historical specimens from both digital and physical herbaria, and computational decoding of taxonomic rationale likely applied by original collectors or appraisers.

**Figure 1.**
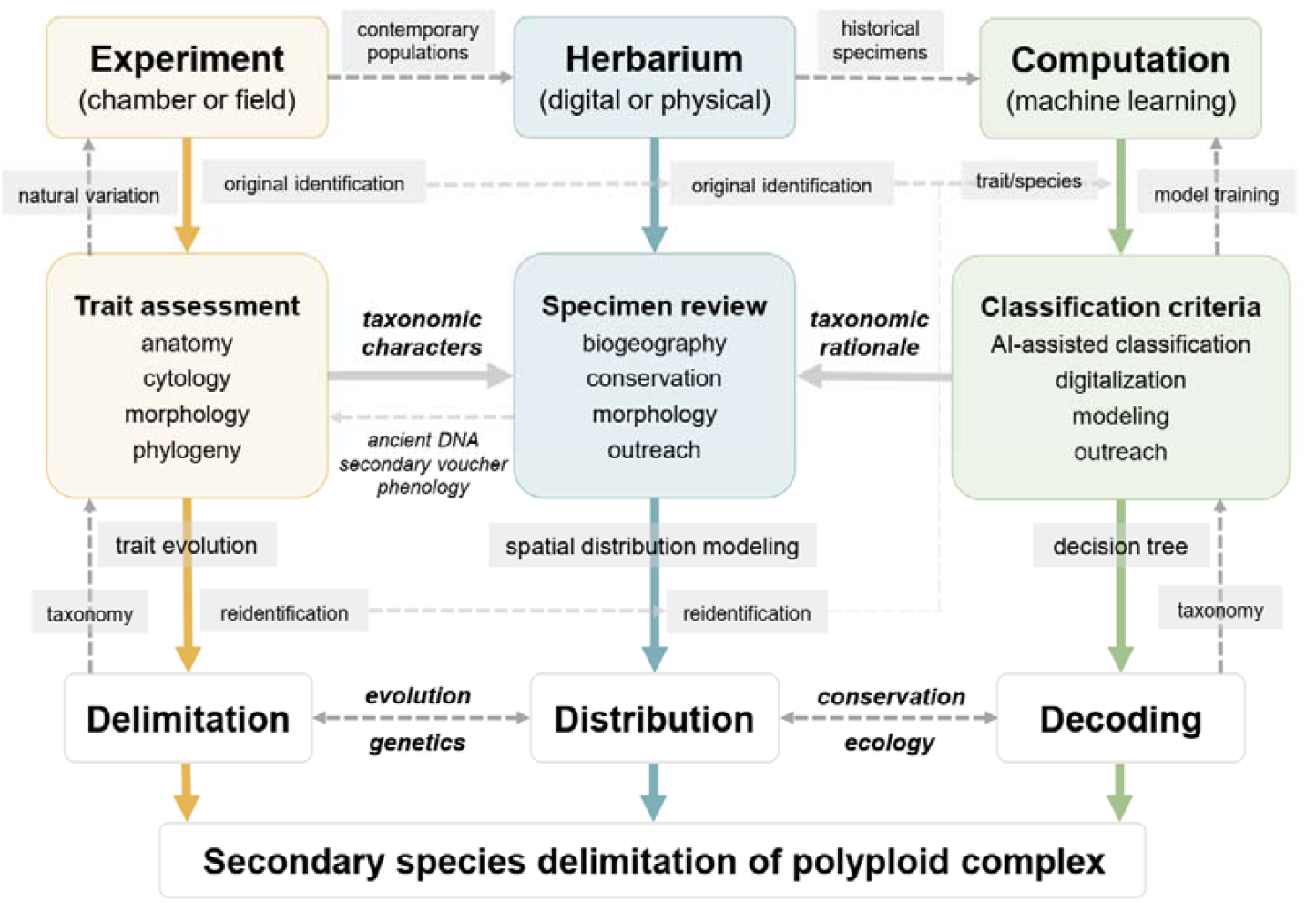
The 3D framework (Delimitation-Distribution-Decoding) for secondary species delimitation in polyploid complexes. The 3D framework integrates three key components: experimental trait analysis for species delimitation, herbarium specimen review (both digital and physical) for reidentification-based distribution modeling, and computational modeling for decoding the classification criteria. In the Delimitation stage, experimental assessments of anatomy, cytology, and morphology are combined with phylogeny-based trait evolution to define species boundaries. In the Distribution stage, stable diagnostic traits are applied to re-identify specimens from herbarium records and field collections, contributing to improved species distribution models. In the Decoding stage, trait and species data are used to train machine learning models (e.g., decision trees) to recover the classification criteria and taxonomic rationale. The decoded rules, corrected SDMs and delimitation not only guide ecological conservation strategies but also enable genetic studies on the evolution of reproductive isolation and the role of trait natural variation in post-speciation adaptation. Together, these components support robust secondary identification, with extension of improving both phenotypic and genetic inference of biodiversity.

The framework has two key features. First, experimental phenotyping under controlled conditions reduces environmental effects and limits trait plasticity, which often obscure true species boundaries in field collections (Hong 2025). This setting enables consistent assessment of both classical diagnostic traits and additional anatomical and cytological features, while allowing for broad and standardized evaluation of diverse accessions. Second, the framework integrates trait data across more than a century of collections, bridging past and present taxonomic perspectives (Burbano & Gutaker 2023). It also incorporates geographically broad sampling and ecological niche modeling, including projections under future climate scenarios. By synthesizing multiple independent lines of evidence, the 3D framework supports more accurate inference of inconspicuous species boundaries in taxonomically complex plant groups (Hörandl 2022; Karbstein et al. 2024).

To demonstrate the utility of this framework, we applied it to a challenging polyploid complex in the mustard family: the *Rorippa dubia–indica* group (Brassicaceae), which exemplifies many of the issues described above. This group includes three cytologically distinct taxa distributed in East Asia: tetraploid *R. dubia* (2*n*=4*x*=32), hexaploid *R. indica* (2*n*=6*x*=48), and the recently described hexaploid *R. hengduanshanensis* (2*n*=6*x*=48) (Zheng et al. 2021). Members of this complex are important crop wild relatives (Castillo-Lorenzo et al. 2024), traditional medicinal herbs (Lin et al. 1995; Hong et al. 2015), and edible wild vegetable in Southern China since the Song Dynasty (960–1279 AD) (Lin 2016). They have also been harnessed in *Brassica* breeding through distant hybridization (Dai et al. 2005; Xin et al. 2024), for their robust resistance to aphids and drought stress (Yamane et al. 1992; Muraoka & Watanabe 1994; Lin et al. 2014; Xu et al. 2016; Sarkar et al. 2017). However, high morphological similarity among taxa in the complex has led to widespread taxonomic confusion.

The group was once treated as a single species (e.g., *R. dubia* was considered as *R. indica* var. *apetala*), with observable variation in traits such as petal number interpreted as intraspecific polymorphism (Gu & Hsu 1986). This taxonomic ambiguity was also reflected in the proliferation of synonyms: over twenty have been reported for *R. dubia* (e.g., *R. heterophylla*) or *R. indica* (e.g., *R. montana*), underscoring long-standing uncertainty in species boundaries (Al-Shehbaz 2001; 2016). As a result, misidentifications have been frequently encountered in both herbarium records and ecological studies (Stuckey 1972; Gu & Hsu 1988; Tu et al. 2019). Users of medicinal plants may inadvertently collect *R. dubia* instead of the more widely used *R. indica*, risking substitution in traditional applications (Lin et al. 1995; Hong et al. 2015). Moreover, members of this complex have expanded their range beyond East Asia, with populations now established in parts of Southeast Asia, North America, and Central America. In these invasive contexts, misidentification may hinder accurate ecological assessment and complicate conservation or management decisions (Rollins 1969; Stuckey 1972).

In this study, we apply the integrative 3D framework to delimit species within the *Rorippa dubia–indica* complex. Specifically, we aim to (1) identify stable morphological traits under controlled conditions; (2) reconstruct population-based phylogenies using uniparentally inherited plastomic variation; (3) use decision tree–based classification models to evaluate the consistency and rationale of species identifications from herbariums; and (4) evaluate the ecological and conservation implications of refined species boundaries. By fusing species delimitation, spatial distribution, and machine learning supervised classification as an iterative system, we clarify species limits within morphologically overlapping polyploid complexes and provide insights for species delimitation in other challenging plant groups.

## Material and method

### Field sampling and phenotyping under growth chamber

From 2017 to 2020, we collected a total of 3136 accessions from 343 natural populations of *R. dubia-indica* complex, initially classified using standard morphological criteria described in regional floras (Al-Shehbaz 2001; 2016). Key morphological traits included cauline leaf shape (sessile or petiolate; margins denticulate or pinnately lobed), presence or absence of petals, fruit curvature (curved or straight), and seed arrangement (uniseriate vs. biseriate) (Fig. S1). To assess seed arrangement, we examined approximately the central 80% of each fruit. The basal and apical ends were excluded from scoring due to the frequent occurrence of irregular or incomplete seed column development in those regions (Gu & Hsu 1986).

**Figure S1.**
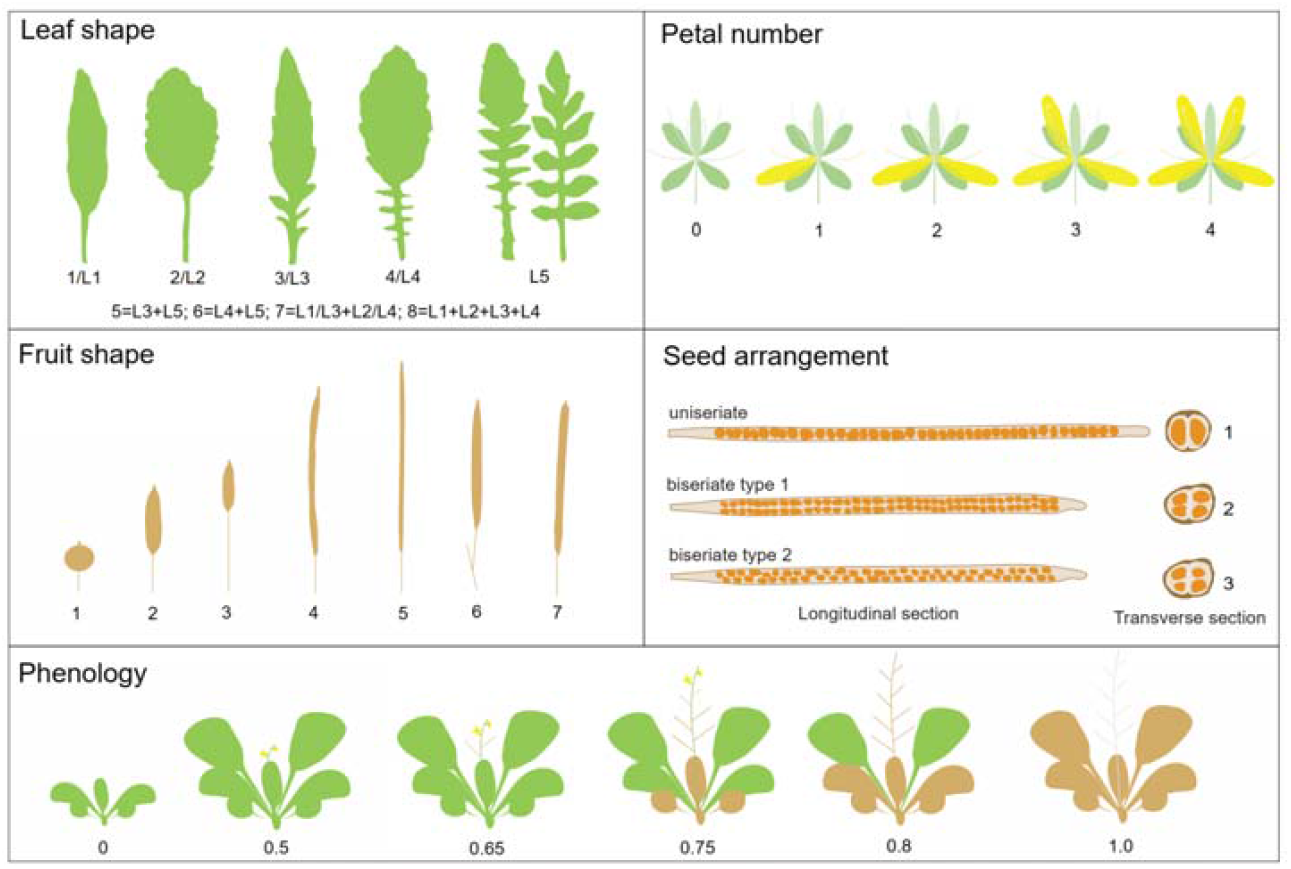
Key morphological traits used for species delimitation in the *Rorippa dubia–indica complex*. Traits include leaf shape types (basic types of L1–L5 and combinations, indexed as 1–8), petal number (0–4), fruit shape (1–7), seed arrangement (1 = uniseriate, 2 = biseriate type 1, 3 = biseriate type 2), and phenology (0–1.0, for specimens).

For secondary assessment, seeds for 686 accessions from 285 populations were germinated and cultivated under controlled conditions in growth chambers in XTBG, Menglun (Fig. S2). Six replicates were used for each accession in randomized 4 × 12 plots. Plants were grown under 16 hours light at 22°C and 60% relative humidity, which was the stable condition for *Rorippa* species. Genome size estimations were performed using flow cytometry on fresh leaves (Zheng et al. 2021). To rigorously examine the natural variation of taxonomic traits (Fig. S1), comprehensive phenotypic evaluations were performed, including petal number recorded from the first five flowers; rosette and cauline leaf shapes indexed systematically from 1 to 8, capturing fundamental morphological variants and their combinations; branching patterns quantified by branch number and length; fruit characteristics assessed by number and length; and seed arrangements classified into three categories: (1) uniseriate, (2) biseriate with parallel seed pairs per valve, and (3) biseriate with imbricate seed pairs per valve.

**Figure S2.**
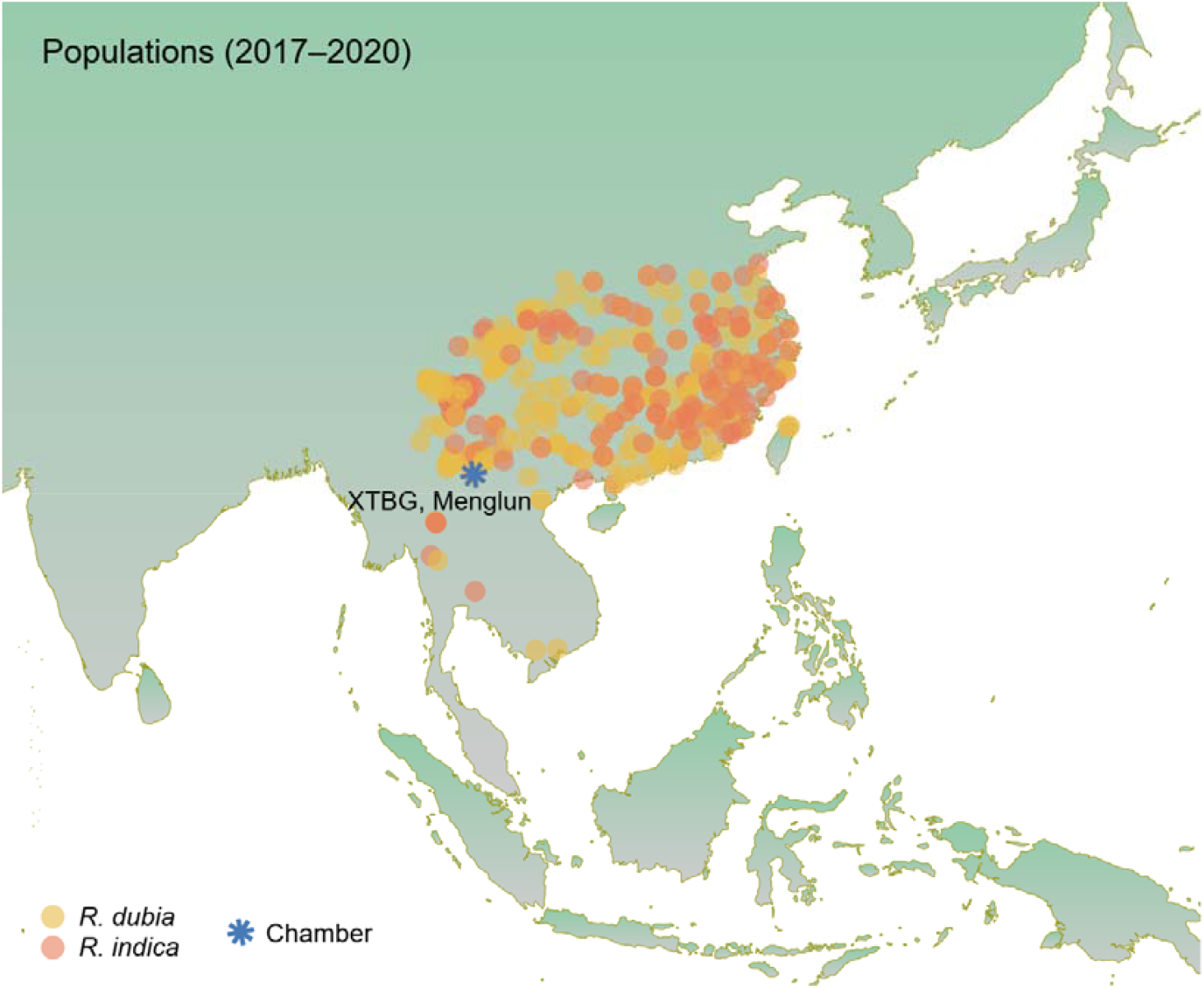
Geographic distribution of *Rorippa dubia and R. indica populati*ons sampled from field collections and grown under experimental conditions. Yellow and orange circles indicate field collection sites for *R. dubia* and *R. indica*, respectively. The blue asterisk marks the location of XTBG (Xishuangbanna Tropical Botanical Garden) in Menglun Town, where accessions were cultivated and phenotyped in a controlled growth chamber.

### Genomic sequencing and phylogenetic analysis

To establish phylogenetic relationships within this polyploid complex, genomic sequencing data were generated and analyzed using published pipeline (Han et al. 2024a). DNAs and libraries were prepared for a total of 161 accessions from experimental examination, comprising 63 accessions of *Rorippa dubia*, 84 of *R. indica*, and 14 of *R. hengduanshanensis*. Half of them were randomly selected and the rest were used to verify species delimitation given the mismatches between original and secondary identifications. For each sample, a minimum of 2 Gb data with paired-end short reads (150 bp) were obtained via the Illumina NovaSeq platform. Published sequences from six other *Rorippa* species and four Brassicale species were used as outgroups. Due to the feasibility of using uniparentally inherited chloroplast genomes for phylogenetic reconstruction, a total of 77 protein-coding genes were extracted from assembled chloroplast genomes and employed in Bayesian phylogenetic analyses using BEAST v.1.8.4 (Drummond et al. 2012). Species delimitation was further validated by applying the Poisson Tree Processes-Maximum Likelihood (PTP-ML) approach (https://species.h-its.org/ptp/) (Zhang et al. 2013).

Phylogenetic signal assessment was performed using Pagel’s λ, where traits with higher λ values indicated greater stability and potential taxonomic utility in the *R. dubia-indica* complex. To identify the most plausible evolutionary pathways between trait pairs, a total of 24 trait-dependent evolutionary scenarios were modeled and evaluated using the Akaike Information Criterion (AIC) (Table S1). These scenarios examined potential dependent or interdependent relationships among phylogenetically delimited taxonomic units, key morphological traits (e.g., seed arrangement, petal number), and cytological characteristics such as ploidy level (tetraploid vs. hexaploid). These analyses were performed using the R package phytools v.2.3-0 (Revell 2012).

**Supplementary Table 1.** Model comparison testing evolutionary relationships among phylogeny, ploidy level, petal number, and seed arrangement. AIC-based model selection results for trait-dependent evolutionary scenarios. The best-supported model (lowest AIC) indicates that ploidy level and phylogeny evolved interdependently (*fit.xy*), followed by models where phylogeny depends on ploidy (*fit.y.x*) or seed arrangement (*fit.y.z*). Bolded models highlight the top three scenarios. AIC weights (AIC.w) and log-likelihood values are provided for evaluating model strength and fit. These results suggest tight co-evolution among phylogeny, genome duplication, and reproductive traits.

| Scenario.ID | Model | AIC | AIC.w | Log-likelihood | Description |
| --- | --- | --- | --- | --- | --- |
| **fit.xy** | **interdependent ploidy-phylogeny** | **52.821** | **0.400** | **-18.410** | **ploidy level and phylogeny evolved interdependently** |
| **fit.y.x** | **dependent phylogeny→ploidy** | **53.821** | **0.243** | **-20.910** | **phylogeny depends on ploidy level, but not the reverse** |
| **fit.y.z** | **dependent phylogeny→seed** | **54.407** | **0.181** | **-21.204** | **phylogeny depends on seed arrangement, but not the reverse** |
| fit.x.y | dependent ploidy→phylogeny | 56.291 | 0.071 | -22.145 | ploidy level depends on phylogeny, but not the reverse |
| fit.z.y | dependent seed→phylogeny | 57.613 | 0.036 | -20.807 | seed arrangement depends on phylogeny, but not the reverse |
| fit.yz | interdependent phylogeny-seed | 57.613 | 0.036 | -20.807 | phylogeny and seed arrangement evolved interdependently |
| fit.x.z | dependent ploidy→seed | 59.489 | 0.014 | -21.745 | ploidy level depends on seed arrangement, but not the reverse |
| fit.xz | interdependent ploidy-seed | 59.489 | 0.014 | -21.745 | ploidy level and seed arrangement evolved interdependently |
| fit.yz | independent phylogeny-seed | 63.091 | 0.002 | -27.545 | phylogeny and seed arrangement evolved independently |
| fit.z.x | dependent seed→ploidy | 64.299 | 0.001 | -26.149 | seed arrangement depends on ploidy level, but not the reverse |
| fit.xy | independent phylogeny-ploidy | 71.466 | 0.000 | -31.733 | phylogeny and ploidy level evolved independently |
| fit.w.z | dependent petal-seed | 80.053 | 0.000 | -32.026 | petal number depends on seed arrangement, but not the reverse |
| fit.z.w | dependent seed→petal | 80.053 | 0.000 | -32.026 | seed arrangement depends on petal number, but not the reverse |
| fit.wz | interdependent petal-seed | 80.053 | 0.000 | -32.026 | petal number and seed arrangement evolved interdependently |
| fit.xz | independent ploidy-seed | 80.528 | 0.000 | -36.264 | ploidy level and seed arrangement evolved independently |
| fit.y.w | dependent phylogeny→petal | 92.179 | 0.000 | -31.733 | phylogeny depends on petal number, but not the reverse |
| fit.wy | interdependent petal-phylogeny | 92.179 | 0.000 | -38.089 | petal number and phylogeny evolved interdependently |
| fit.x.w | dependent ploidy→petal | 92.602 | 0.000 | -38.301 | ploidy level depends on petal number, but not the reverse |
| fit.wx | interdependent petal-ploidy | 92.602 | 0.000 | -38.301 | petal number and ploidy level evolved interdependently |
| fit.w.y | dependent petal→phylogeny | 95.205 | 0.000 | -41.602 | petal number depends on phylogeny, but not the reverse |
| fit.w.x | dependent petal→ploidy | 96.348 | 0.000 | -42.174 | petal number depends on ploidy level, but not the reverse |
| fit.wy | independent petal-phylogeny | 97.161 | 0.000 | -44.581 | petal number and phylogeny evolved independently |
| fit.wz | independent petal-seed | 106.223 | 0.000 | -49.112 | petal number and seed arrangement evolved independently |
| fit.wx | independent petal-ploidy | 114.598 | 0.000 | -53.299 | petal number and ploidy level evolved independently |

### Specimen collection and identification

Specimens of *R. dubia* and *R. indica* were collected across their known distributions in Eastern and Southern Asia (Fig. S3). Initial identifications were performed using photographic records through the Chinese Virtual Herbarium (CVH, https://www.cvh.ac.cn/; last accessed on August 18, 2023), and subsequently verified through direct examination of specimens housed in five major Chinese herbaria (IBSC, KUN, NAS, PE, and WUK; last accessed on October 12, 2023). A total of 2,015 specimens were reviewed, including 573 initially labeled as *R. dubia* (28.437%) and 1,418 as *R. indica* (70.372%) during collections from 1893 to 2021. Of these, 1,811 specimens—526 *R. dubia* and 1,251 *R. indica*—were confirmed through both virtual and physical examination. Approximately 6.153% and 3.970% of specimens were accessible only via herbarium visits or CVH images, respectively.

**Figure S3.**
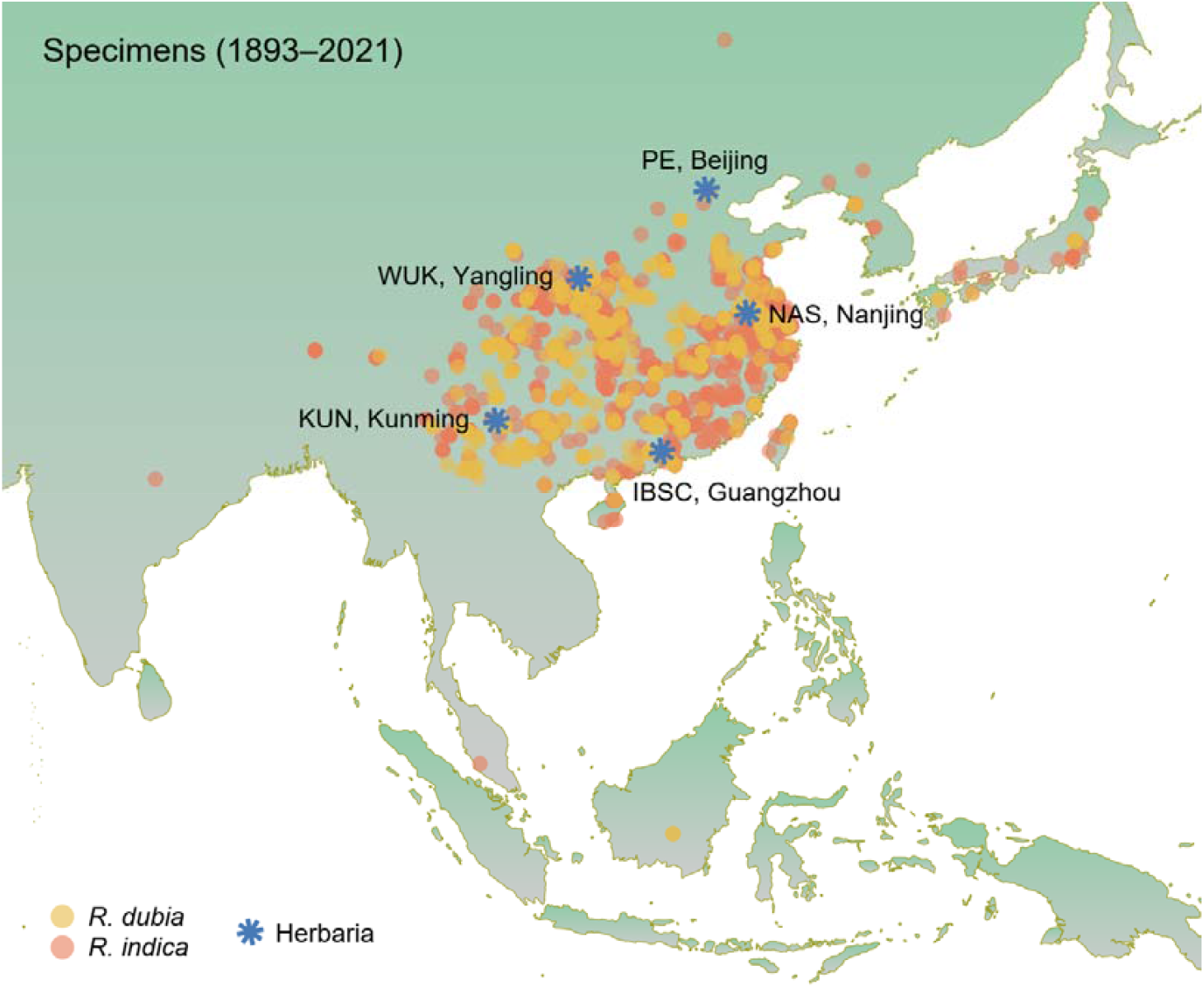
Geographic distribution of *Rorippa dubia and R. indica* specimens and five major herbaria used for taxonomic review. Specimen localities are shown across Eastern and Southeastern Asia, with yellow and orange points representing *R. dubia* and *R. indica*, respectively. Blue asterisks indicate the locations of five key herbaria where physical specimens were examined: IBSC (Guangzhou), KUN (Kunming), NAS (Nanjing), PE (Beijing), and WUK (Yangling).

Reidentifications were conducted using a standardized set of morphological traits (Fig. S1), including leaf shape (indexed categorically), petal number, fruit morphology (coded on a 1–7 scale from globose to linear), seed arrangement (uniseriate or biseriate types), and phenological status. Phenological stages were scored numerically from 0 (seedling) to 1 (fully senesced), following the index developed by Wu & Colautti (2022). Plants of *R. hengduanshanensis* were included only during the reidentification phase, as no specimens were available prior to its formal description (Zheng et al. 2021).

To check the accuracy of specimen-based reidentification, a total of 15 specimens were randomly selected for whole-genomic sequencing, including 6 samples initially wrongly identified as *R. dubia* and 9 as *R. indica*. The same pipeline was implied for chloroplast genomic assembly and annotation. The tested samples were placed into the former plastome tree using phylogenetic placement (Czech et al. 2020).

### Decision tree of species identification

To evaluate the phenotypic and metadata variables influencing taxonomic misidentification in the *R. dubia-indica* complex, we constructed decision tree models using both conditional inference trees and classification and regression trees (CART). Analyses were performed using the R packages party v.1.3-15 for inference trees and rpart v.4.1-10 for CART (Hothorn et al. 2006).

Four datasets were assembled: two excluding entries with missing data for conditional inference tree construction (1053 and 1003 entries for original or secondary identification), and another two retaining all records for CART modeling (1900 and 1793 entries). Each dataset was randomly partitioned into training (70%) and testing (30%) sets. The model formula used was: *y* ∼ Leaf + Petal + Fruit + Seed + Phenology, where *y* represents the binary classification of species assignment and the explanatory variables include leaf shape (Leaf), petal number (Petal), fruit morphology (Fruit), seed arrangement (Seed), and phenological state (Phenology). To reduce bias due to features with extraordinary unique values, taxonomic traits were all scored when incorporated in the model (Fig. S1).

For the CART models, we specified a minimum split size (minsplit) of 50 and a minimum number of observations in any terminal node (minbucket) of 20. The complexity parameter (cp) was tuned by minimizing cross-validation error (xerror). To clarify the stability and overfitting, CART models were replicated 100 times to identify the best-performing tree structure. Model performance was assessed using receiver operating characteristic (ROC) curves and the area under the ROC curve (AUC). Confusion matrices were generated using the caret package v.6.0-90 to evaluate classification accuracy and misclassification patterns. From these models, decision rules were extracted to identify key traits influencing species assignment and to reveal systematic errors in specimen identification.

### Ecological effects of species delimitation

The ecological consequences of species misidentification were systematically assessed through our developed pipeline under the 3D framework (Fig. 1). Particularly, kernel density estimations and species distribution modeling (SDM) were integrated to explore spatial distribution patterns between original and secondary identifications. We focused on *R. dubia* and *R. indica* for ecological analyses, as *R. hengduanshanensis* had not been recognized as a distinct species in earlier specimen records (Zheng et al. 2021).

### □. Kernel density of specimen distribution

To evaluate the geographic distribution and density shifts of *R. dubia* and *R. indica* before and after reidentification, kernel density analysis was conducted using ArcMap v.10.8. Only specimens with unique GPS coordinates and confirmed through artificial examination were included. A total of 1,065 occurrence records were retained for *R. dubia*, consisting of 470 records prior to re-identification and 595 afterward. For *R. indica*, 1,255 records were analyzed—717 pre-identifications and 538 post-identifications. All GPS data were projected onto a base map of China. Kernel density surfaces were generated using a 2-mile search radius and a standardized cell size determined by the highest-resolution dataset. Raster masks were applied to constrain analysis to the relevant geographic extent. Spatial patterns were visualized to identify shifts in distribution associated with reidentification and assess potential herbarium-related geographic biases.

To quantify spatial differentiation between original and re-identified occurrences for *R. dubia* and *R. indica*, niche overlap analyses were conducted based on their modeled distributions. Species occurrence data were aggregated by Chinese municipal administrative units using spatial join in ArcGIS. Species abundances per unit were compiled into a species-by-site matrix. Niche overlap indices, including Czechanowski’s, Levins’s, simplified Morisita’s, Petraitis’, Pianka’s and Schoener’s metrics, were calculated using the R package spaa v.0.2.5 (https://github.com/helixcn/spaa). Multiple *T*-tests were performed to evaluate whether observed overlaps significantly differed from random expectations.

### □. Environmental data and occurrence coordinate

Nineteen bioclimatic variables at 2.5 arc-minute resolution were obtained from the WorldClim database (https://www.worldclim.org/). These variables were extracted at each specimen occurrence point using the raster v.3.6-23 package in R. Pearson correlation analysis was used to identify and remove highly correlated variables (|*r*| ≥ 0.8), thereby minimizing overfitting in subsequent ecological niche models. The most informative and uncorrelated seven environmental variables were retained for further modeling, including bio1, bio2, bio3, bio7, bio8, bio12, and bio15.

Geographic coordinates from original collections are typically unevenly distributed due to collection bias, which can distort distribution models if uncorrected. To reduce sampling bias and spatial autocorrelation in species distribution modeling (SDM), specimen occurrence data were spatially thinned using the R package spThin v.0.2.0 (Aiello-Lammens et al. 2015), with thinning distances ranging from 0 to 150 km at 5-km increments. For each thinned dataset, the Average Nearest Neighbor Index (ANNI) was calculated in ArcGIS to assess spatial randomness. Three datasets per species with ANNI values closest to 1 were selected as spatially representative for downstream modeling.

Each dataset was partitioned into a training set (75%) and a test set (25%). SDMs were constructed under 6 × 13 model combinations, incorporating six feature class settings (H, L, LQ, LQH, LQHP, LQHPT) and thirteen regularization multipliers (rm = 0.1–6.0, in 0.5 increments) using the R package ENMeval v.2.0.4 (Kass et al. 2021). Model performance was evaluated using corrected Akaike Information Criterion (AICc; lower is better) and the area under the receiver operating characteristic curve (AUC; higher is better, based on test data). The optimal model and thinning distance combinations were as follows: *R. dubia* (original): 65 km thinning, ANNI = 0.983, AUC = 0.897, model = LQ + rm2.5; *R. indica* (original): 90 km, 1.011, 0.866, LQ + rm1.0; *R. dubia* (reidentified): 70 km, 0.975, 0.901, LQ + rm0.1; *R. indica* (reidentified): 75 km, 0.989, 0.881, LQHPT + rm3.0.

### □. Ecological niche modeling

Species distribution models (SDMs) were developed using Maximum Entropy (MaxEnt) modeling, implemented through the R package ENMeval v.2.0.4 (Kass et al. 2021). Models combined latitude–longitude coordinates of thinned occurrence records with the subset of selected, uncorrelated seven bioclimatic variables. SDMs were run under the optimal thinning distances, feature class (fc) combinations, and regularization multipliers (rm) identified previously using the checkerboard2 partitioning scheme.

To evaluate the potential impacts of climate change on species distributions, future climate projections were obtained from the Coupled Model Intercomparison Project Phase 5 (CMIP5) dataset under three representative concentration pathways (RCPs) of greenhouse gas: RCP4.5 (low emissions), RCP6.0 (medium emissions), and RCP8.5 (high emissions). Three general circulation models (GCMs)—MIROC5, NorESM1-M, and GISS-E2-R—were used to capture a range of atmospheric and spatial resolutions. Future SDMs were constructed using the same MaxEnt settings as the present-day models. Predicted suitability layers from the three GCMs were averaged for each RCP scenario to generate ensemble forecasts of future habitat distributions for both *R. dubia* and *R. indica*.

## Results

### Delimitation of taxonomic character in Rorippa complex

Morphological and cytological traits were examined from chamber-grown individuals to assess their diagnostic potential while minimizing environmental variation within the *Rorippa dubia–indica* complex (Fig. S4). Several traits displayed discrete or bimodal distributions, suggesting interspecific differentiation. Notably, seed arrangement, genome size and petal number, showed sharply separated peaks corresponding to known cytotypes (Shapiro– Wilk tests, *W* ≤ 0.686), reinforcing their utility for species delimitation (Fig. S4). Traits such as leaf shape, plant height, and fruit length exhibited continuous distributions (*W* ≥ 0.828), consistent with higher levels of phenotypic plasticity. Together, these findings support the potential use of stable and discretely distributed traits for reliable identification within the *R. dubia–indica* complex.

**Figure S4.**
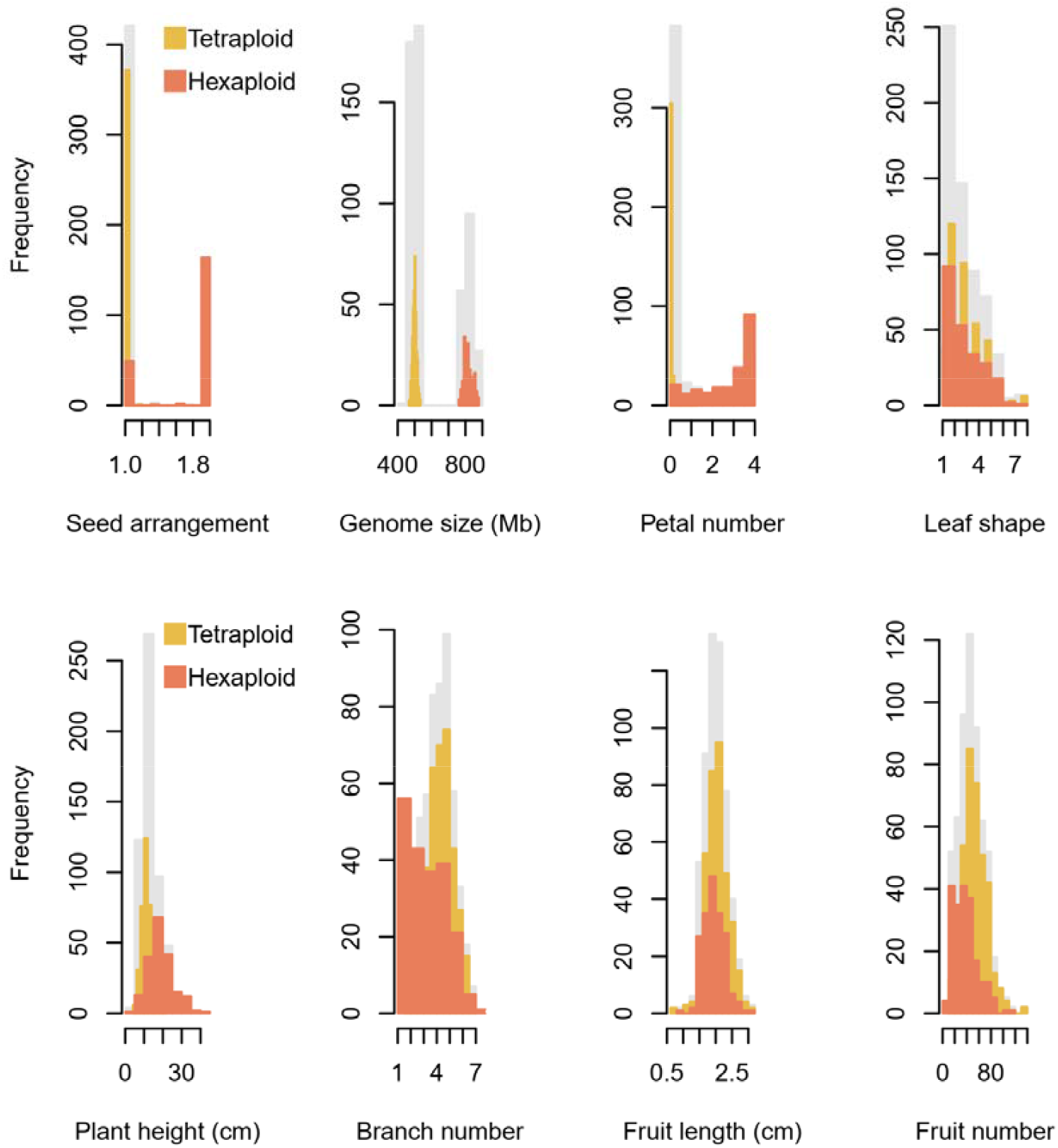
Trait distributions of *Rorippa* accessions under growth chamber. Histograms show the frequency distributions of morphological and cytological traits assessed in chamber-grown plants (grey), grouped by ploidy level: tetraploid (yellow) and hexaploid (pink) accessions. Traits include seed arrangement, leaf shape, genome size, petal number (mean of first five flowers), fruit length, plant height, branch number, and fruit number.

A well-resolved chloroplast genome-based phylogeny provided clear support for species relationships within the *R. dubia–indica* complex (Fig. 2). Among the morphological and cytological traits examined, seed arrangement, petal number, and genome size (or ploidy level) exhibited the strongest phylogenetic signals, as indicated by high Pagel’s λ values (Table 1). Importantly, the most phylogenetically informative traits also demonstrated clear interspecific differentiation, reinforcing their taxonomic utility (Fig. 2b). Plants of *R. dubia* were uniseriate, tetraploids, and typically lacking petals. In contrast, *R. indica* plants were biseriate, hexaploid, and rarely petal-less. Plants of *R. hengduanshanensis* were hexaploid with four petals, but consistently displayed uniseriate. Minor inconsistencies were observed between phylogeny-based and trait-based classifications. Specifically, three hexaploid accessions (proportion = 1.8%) were embedded within the *R. dubia* clade based on chloroplast phylogeny, and two accessions exhibited variation in seed arrangement among replicates (1.2%), congruence with prior report (Gu & Hsu 1986). Overall, the screened traits are evolutionarily conserved and thus highly informative for species delimitation. In contrast, traits such as leaf shape and fruit length showed low phylogenetic signal, likely reflecting greater phenotypic plasticity or convergent evolution.

**Table 1.** Phylogenetic signal of taxonomic traits estimated by Pagel’s λ. Traits are ranked by their degree of phylogenetic signal (*λ*), indicating the extent to which trait variation is correlated with phylogeny. Seed arrangement, petal number, and genome size exhibit the strongest phylogenetic signals, suggesting they are evolutionarily conserved and informative for species delimitation. Lower *λ* values for traits like leaf shape and fruit length reflect greater phenotypic plasticity or lability. *P*-values indicate the statistical significance of each signal against the null hypothesis (i.e., *λ* = 1.000) through likelihood ratio tests.

| Trait | Pagel's $\lambda$ | *P*-value |
| --- | --- | --- |
| Seed arrangement | 0.924 | 1.680E-03 |
| Petal number | 0.923 | 2.111E-03 |
| Genome size | 0.794 | 4.272E-09 |
| Plant height | 0.357 | 1.402E-15 |
| Fruit number | 0.280 | 6.986E-14 |
| Branch number | 0.238 | 7.665E-16 |
| Fruits length | 0.114 | 1.064E-20 |
| Leaf shape | 0.034 | 1.612E-24 |

**Figure 2.**
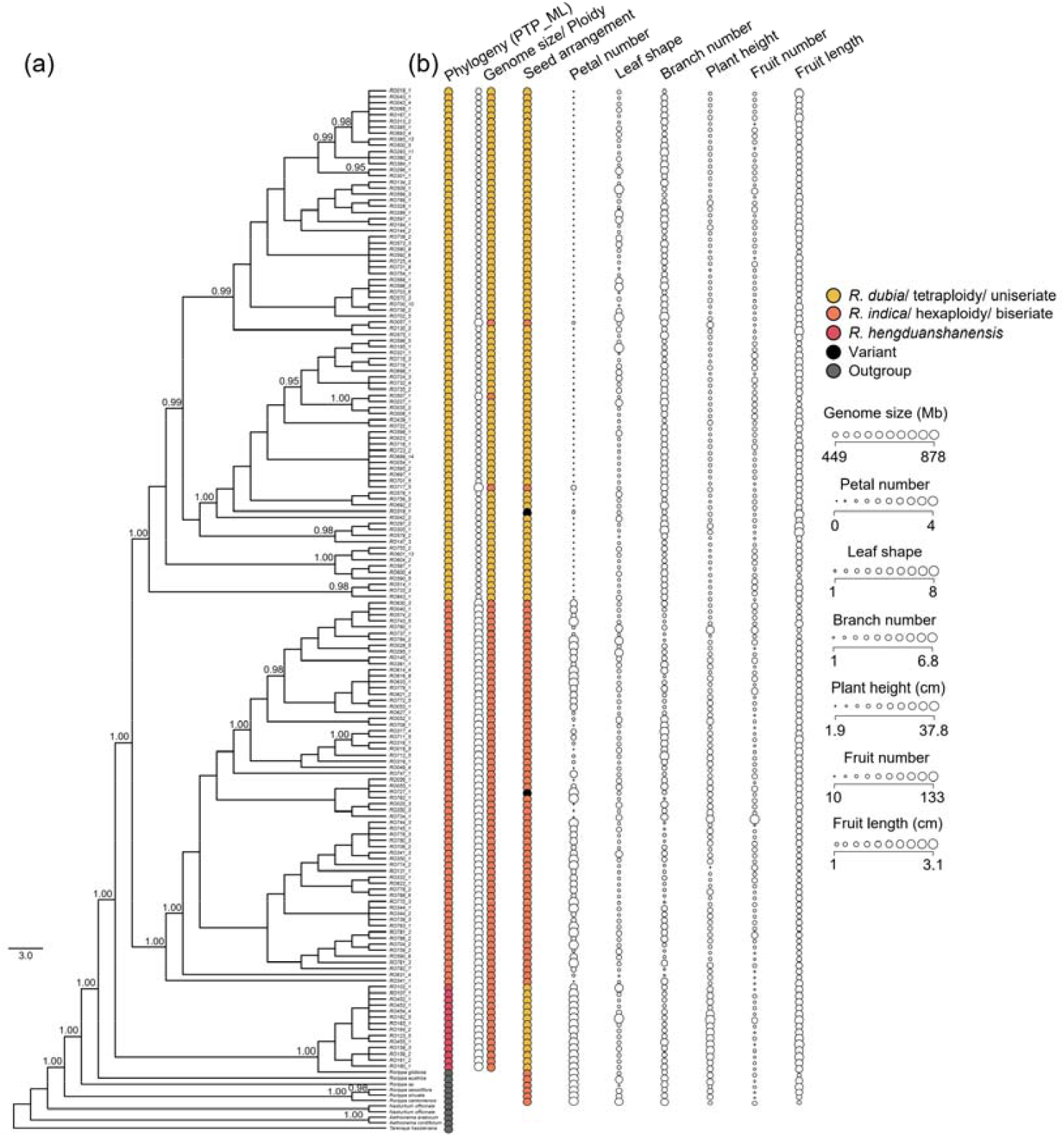
Integrative species delimitation in the *Rorippa dubia–indica* complex. (a) Bayesian phylogeny based on chloroplast genome sequences, showing relationships among accessions and outgroups. Posterior probabilities >0.9 are indicated at key nodes, with scale bar showing branch length. (b) Overview of delimitation criteria integrating phylogeny (PTP-ML), cytological data (genome size and ploidy level), and phenotypic traits (seed arrangement, petal number, leaf shape, branch number, plant height, fruit number, and fruit length). Trait categories include both qualitative (indicated by colorized pies, e.g., seed arrangement) and quantitative metrics (indicated by open pies with varying sizes, e.g., genome size, fruit length).

Trait interaction analyses further revealed evolutionary coupling among characters (Table S1). Model comparisons based on trait-dependent evolutionary scenarios supported an interdependent evolution of ploidy level and phylogeny (weight of AIC or AIC.w = 0.400), with additional support for models in which phylogeny depends on either ploidy or seed arrangement (AIC.w = 0.243 or 0.181). These results suggest traits, such as ploidy level and seed arrangement, are critical taxonomic characters related with speciation.

### Species identification

Enhanced identification accuracy was achieved through direct re-examinations of specimens at five major herbaria and double-checked by plastomic classification (Figs. 2 & 3). Substantial misidentification rates were observed across different data sources, with additional taxonomic transitions involving *R. hengduanshanensis* and other *Rorippa* species (Fig. 3b). For instance, reidentification based on Chinese Virtual Herbarium (CVH) images showed a high misidentification rate of 49.7% for *R. indica* incorrectly labeled as *R. dubia* (Fig. 3c). This rate was reduced to 36.3% after direct examination of physical specimens during herbarium visits (Fig. 3d). Even field-based identifications by experienced taxonomists exhibited considerable error, with a misidentification rate of 31.5% (Fig. 3e).

**Figure 3.**
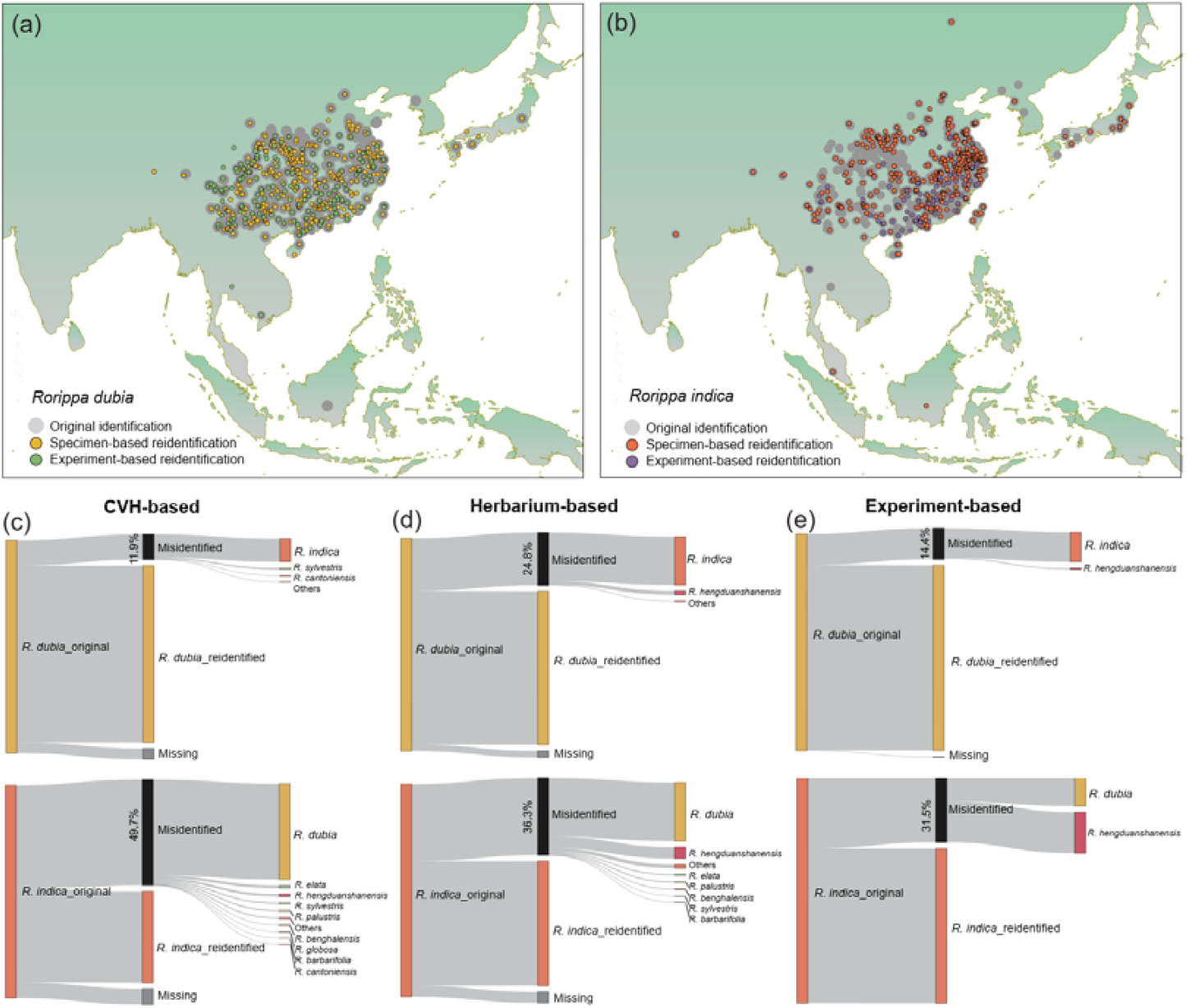
Geographic distribution and reidentification of *Rorippa dubia* and *Rorippa indica* accessions. (a−b) Sampling localities for *R. dubia* (a) and *R. indica* (b) across Asia. Points are colored by identification status: gray = original identification, yellow/red = reidentified based on herbarium specimens, green/purple = reidentified through experimental cultivation. The maps highlight extensive geographic overlap and reveal notable shifts in species assignment following reassessment. (c−e) Flow diagrams from left to right depict changes in species delimitation from original to reidentification for CVH-based virtual specimens (c), herbarium specimens (d), and experimentally grown plants (e). Line thickness represents the proportion of specimens reclassified (black), confirmed (yellow for *R. dubia*, pink for *R. indica*), or missing (grey). Notable misidentification rates are labeled for *R. dubia* and *R. indica*, with additional transitions involving *R. hengduanshanensis* and other *Rorippa* species.

On the side of *R. dubia*, misidentification was also prevalent—24.8% for herbarium-based specimens—often due to subtle variation in petal presence or seed arrangement not visible in initial observations (Fig. 3d). Misidentification rates from CVH-based and experiment-based identifications were relatively lower but still significant, at 11.9% and 14.4%, respectively. These findings underscore the challenges of accurate species delimitation in morphologically continuous polyploid complexes and highlight the need for integrated taxonomic approaches.

### Trait use and identification rationale

Decision tree models based on morphological and phenological traits revealed clear differences in classification criteria and performance between original and secondary identification (Figs. 4 & S5). The structure of the trees varied notably in complexity. Initial identifications often relied heavily on plastic traits—particularly leaf and fruit shape—resulting in deeper branching patterns and reduced classification accuracy (Fig. 4a). In contrast, reidentification incorporated a broader suite of traits, including seed arrangement and petal number, which led to shallower, more streamlined trees with improved discriminatory power (Fig. 4b).

**Figure 4.**
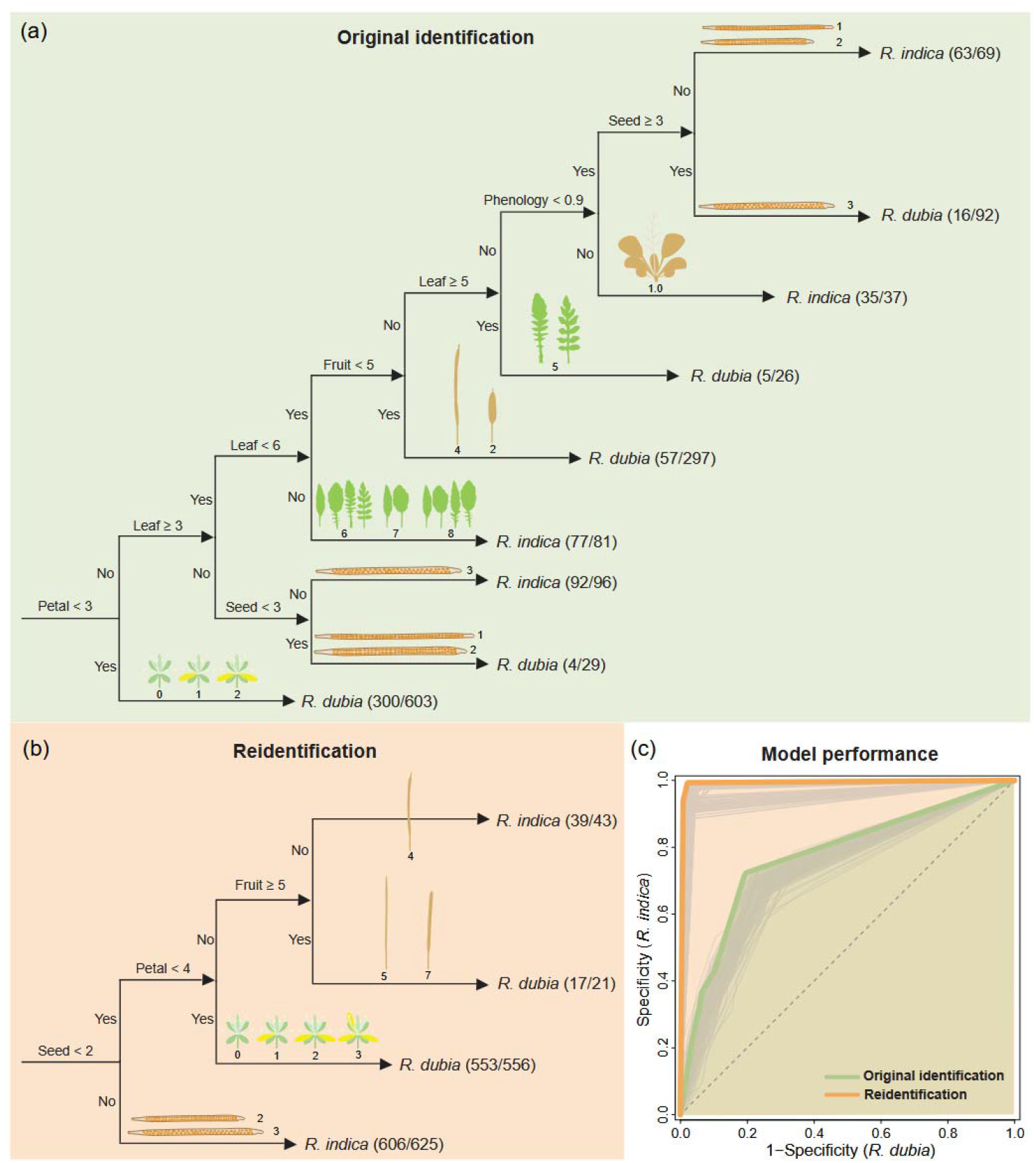
Decision tree models for species delimitation of *Rorippa dubia* and *Rorippa indica*. (a) and (b) show the best-performing conditional inference trees based on 100 simulation replicates for the original identification (a) and reidentification (b), constructed using morphological and phenological traits. Each node represents a decision rule based on trait thresholds, with terminal nodes indicating species assignment and sample counts. (c) Performance evaluation across simulations, shown as receiver operating characteristic (ROC) curves for the 100 simulation replicates (light grey), and highlighted the best-performing model with the highest value of area under ROC curve (AUC) for the original identification (green) and reidentification (orange). Higher AUC values in the reassessment reflect improved identification consistency. The trees highlight differences in trait usage and diagnostic efficiency between rounds, with original identification models relying more on plastic traits and second identification models favoring integrative, phylogenetically informative traits.

**Figure S5.**
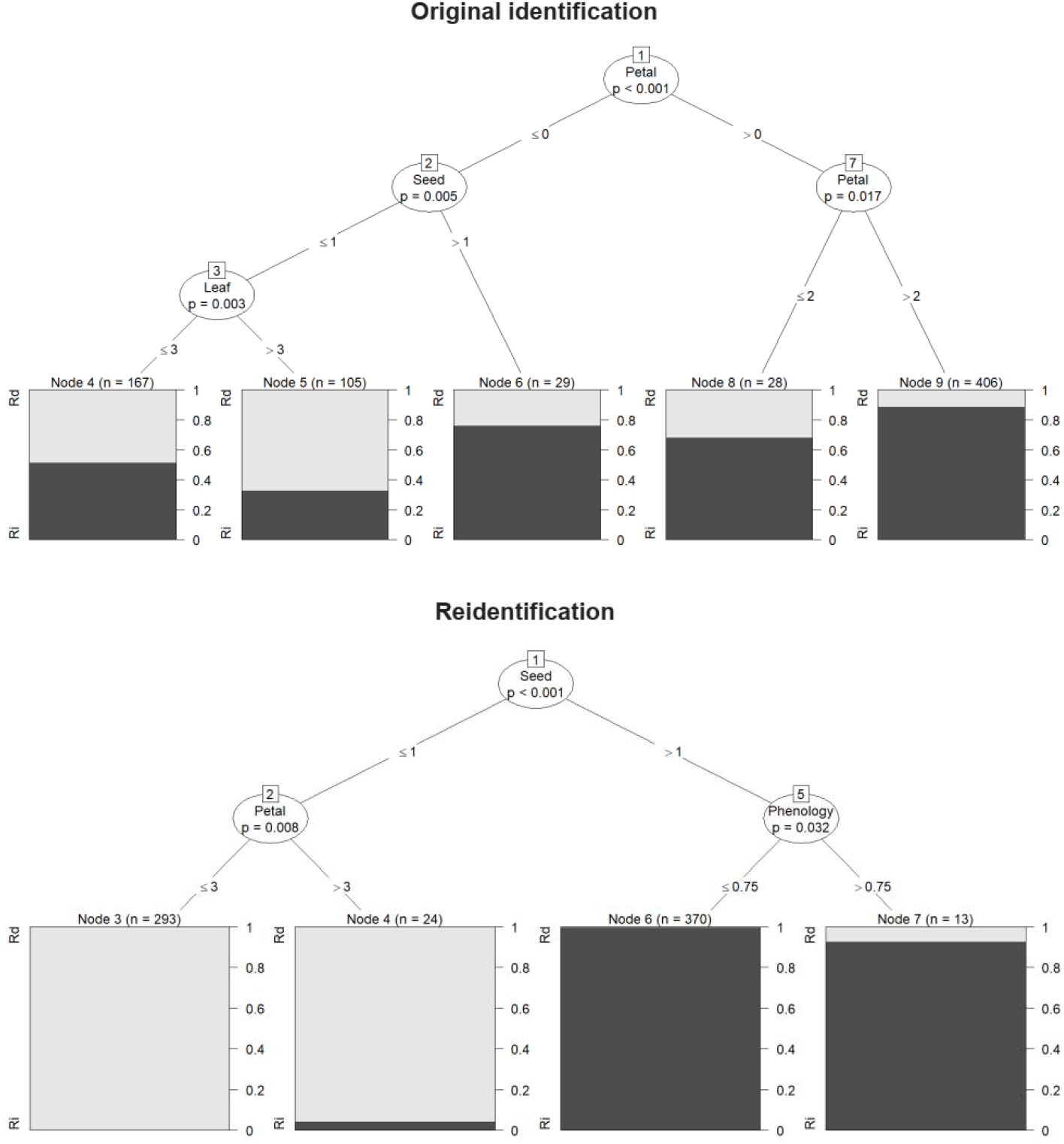
Conditional inference trees for species delimitation of *Rorippa dubia* and *Rorippa indica*. Decision trees constructed using morphological and phenological traits show classification logic for original identifications (above) and reidentifications (below). Each split is based on statistically significant trait thresholds, with terminal nodes indicating species assignment and classification accuracy for *R. dubia* (abbreviated by Rd, light grey) and *R. indica* (Ri, dark grey). The trees highlight how trait importance and decision structure shifted between rounds, with reidentification relying more on stable, phylogenetically informative traits and resulting in more consistent outcomes.

These differences in tree structure and performance metrics reflect the consequences of trait choice: reliance on variable traits increased ambiguity and required more decision steps, whereas the inclusion of evolutionarily conserved traits enabled clearer, more consistent classifications. This pattern was further supported by model performance: reidentification trees exhibited higher predictive accuracy (AUC = 0.988) and greater specificity (0.984) compared to the original models (0.775 and 0.803) (Fig. 4c).

Together, these findings indicate that the depth and topology of decision trees offer insight into the diagnostic value of individual traits. Specifically, the number of decision steps and the nature of trait splits serve as proxies for taxonomic complexity and classification reliability in morphologically overlapping groups.

### Ecological effect of species misidentification

Reassessment of species boundaries revealed extensive geographic shifts within or between *R. dubia* and *R. indica*, accompanied by notable shifts in species assignments following taxonomic reidentification (Figs. 5 & S6). The results indicated what are their actual distribution areas after taxonomic refinement. For *R. dubia*, the revised distribution became more cohesive and spatially concentrated, in contrast to the previously fragmented and scattered pattern. In *R. indica*, reidentification resulted in a clearer separation of populations across central and southern China, suggesting increased spatial isolation (Fig. S6; niche overlap indices, mean ± SD = 0.814 ± 0.066; one-sample *T*-test, *P* = 0.001) and improved resolution of species-specific occurrence data. These shifts emphasize how taxonomic refinement can substantially alter our understanding of species’ spatial distribution and biogeographic structure.

**Figure 5.**
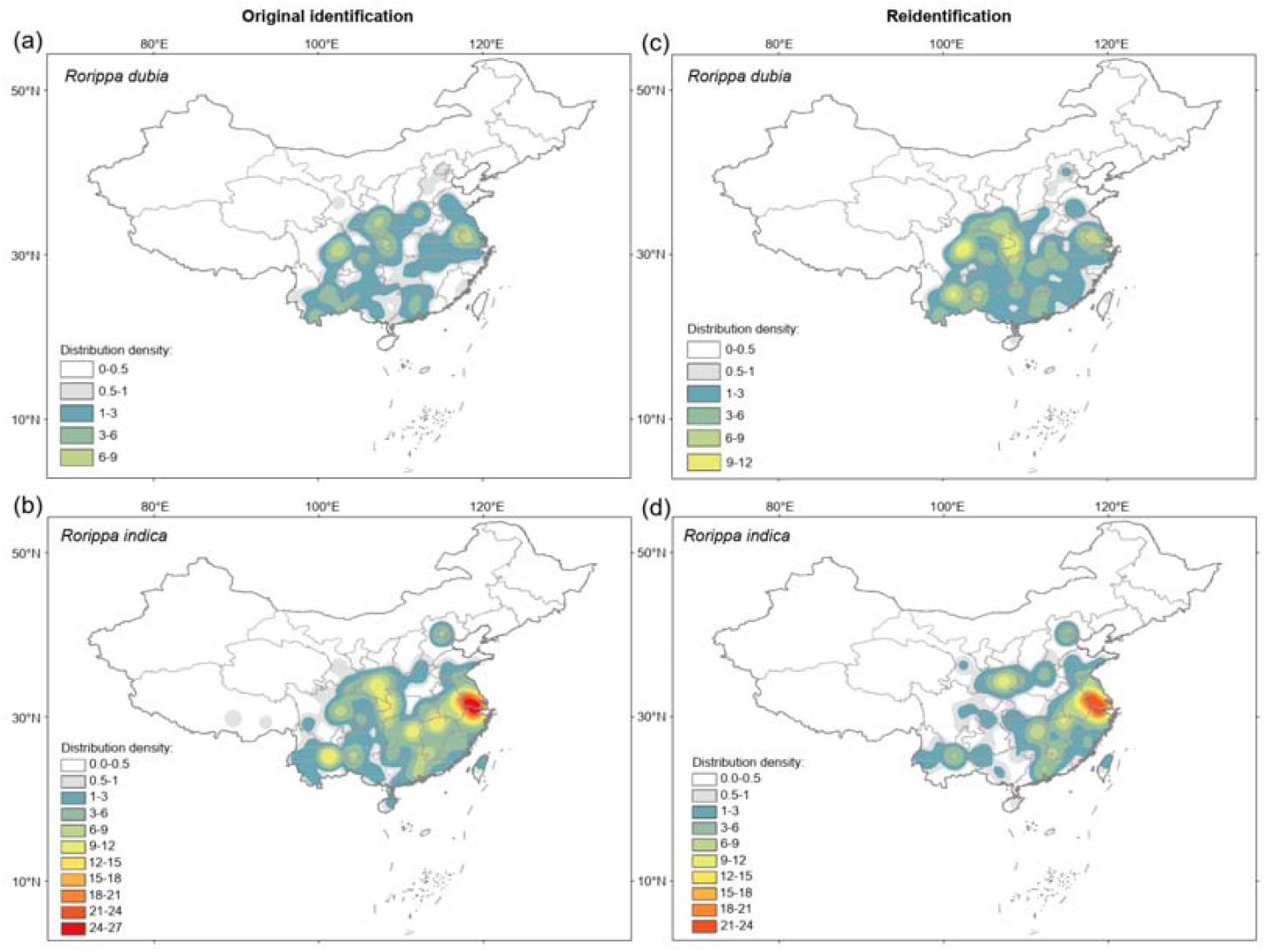
Changes in distribution density before and after reidentification of *Rorippa dubia* and Rorippa *indica*. Kernel density estimates of specimen occurrences across China for *R. dubia* (a−c) and *R. indica* (b−d), shown before (a−b) and after (c−d) taxonomic reidentification. Warmer colors indicate higher local specimen density. The maps reveal significant geographic shifts in species assignment, underscoring the ecological implications of misidentification.

**Figure S6.**
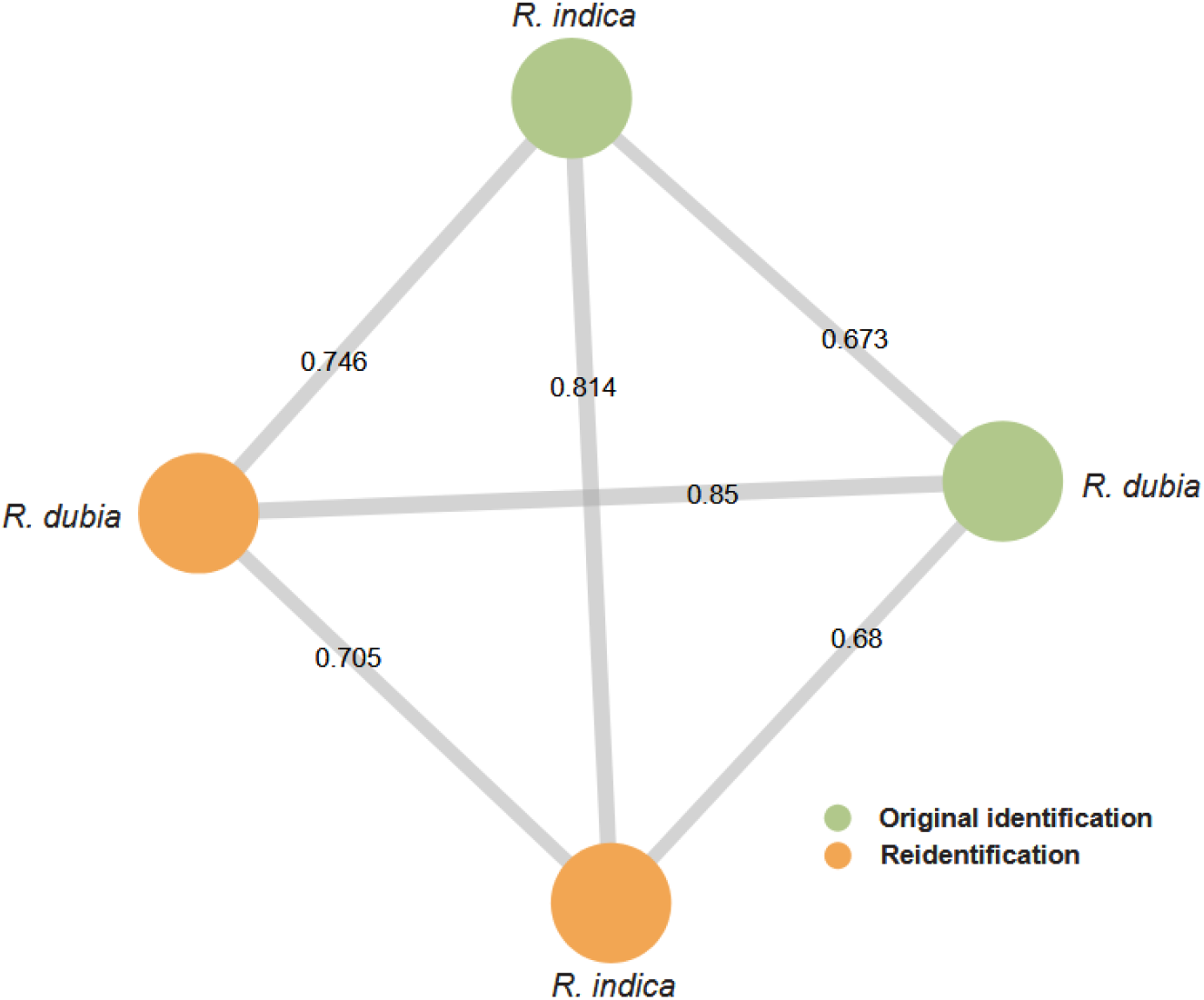
Network of pairwise ecological niche overlap before and after species reidentification. Nodes represent *Rorippa dubia* and *R. indica* under original (green) and reidentified (orange) classifications. Edge thickness corresponds to the degree of niche overlap (mean value across six indices), with values labeled on each connecting edge. Niche overlap was calculated using species distribution models and occurrence data, illustrating how reidentification affects spatial niche estimates. The network reveals moderate to high overlap among original and secondary identities, with notably higher congruence between reidentified taxa (e.g., *R. dubia*–*R. indica*, 0.705 vs. 0.673). This suggests taxonomic refinement alters the inferred ecological niches of cryptic taxa.

Species distribution models (SDMs) constructed before and after reidentification revealed substantial changes in predicted ecological niches for *R. dubia* and *R. indica*, reflecting enhanced species differentiation following taxonomic refinement (Fig. 6). However, the impact of reidentification on niche predictions differed markedly between the two species. For *R. dubia*, niche models showed a contraction of suitable habitat (probability > 0.8), particularly with the loss of predicted presence around the middle and lower reaches of the Yangtze River along the latitude 30°N (Fig. 6a). In contrast, *R. indica* exhibited a notable expansion of predicted niche space across central and southeastern China, although some peripheral areas in the western part of its former range were no longer well-fitted (Fig. 6b). These contrasting patterns highlight how taxonomic precision can reshape ecological interpretations and inform species-specific conservation strategies.

**Figure 6.**
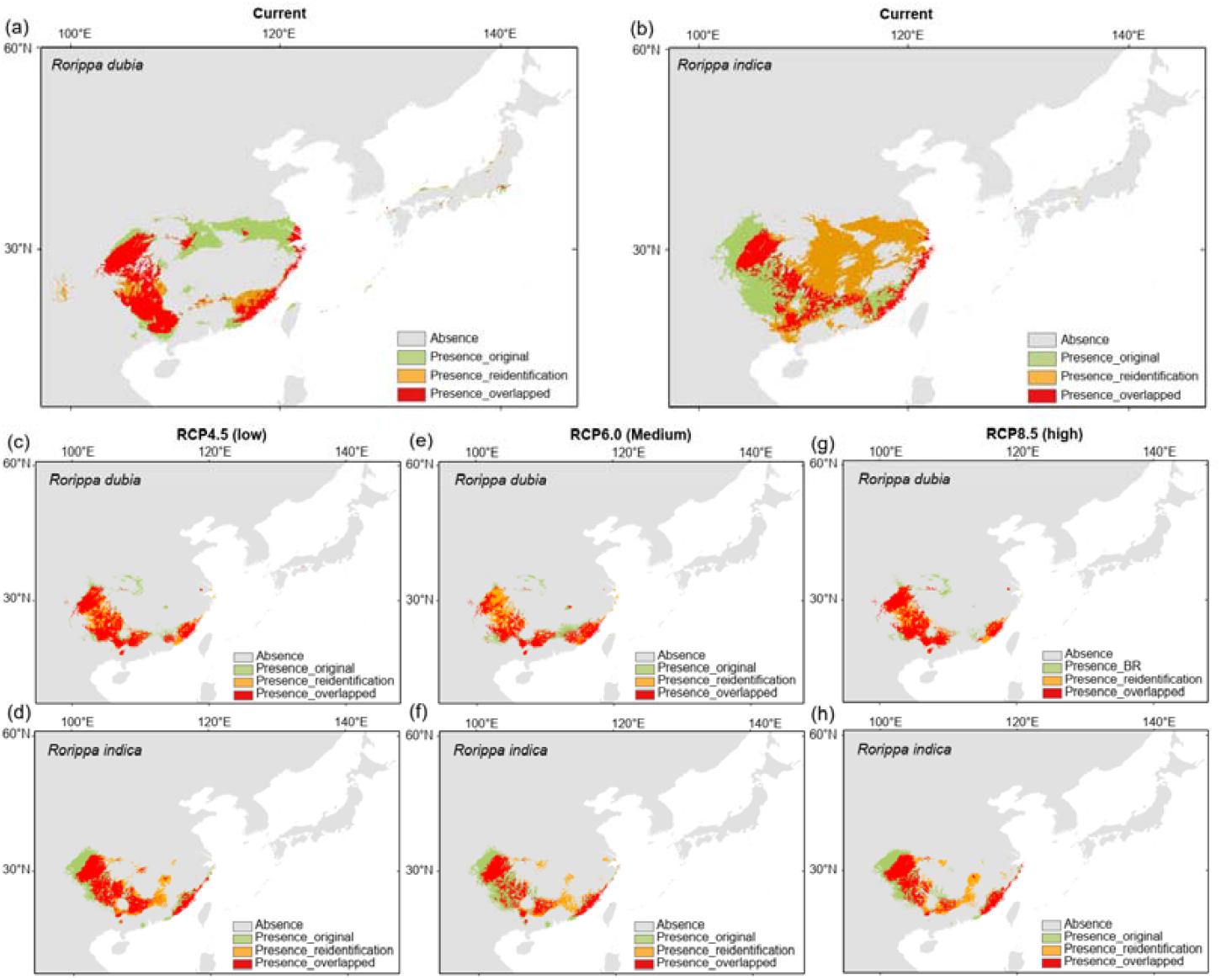
Predicted ecological niches of *Rorippa dubia* and *Rorippa* indica before and after species reidentification. Species distribution models (SDMs) show areas of high habitat suitability (probability ≥ 0.8) for *R. dubia* and *R. indica* under current (a–b) and future climate conditions, including RCP4.5 (low emissions; c–d), RCP6.0 (medium emissions; e– f), and RCP8.5 (high emissions; g–h) scenarios. Colored regions indicate habitat predicted only for original identification (green), only for reidentification (orange), and areas of overlap between the two (red). Gray areas represent unsuitable habitats (probability < 0.8). The projections highlight the differing impacts of taxonomic refinement and climate change on habitat availability for the two species, with *R. indica* showing greater niche shifts and vulnerability under future scenarios.

Species distribution models projected under three representative concentration pathways of future greenhouse gas (RCP4.5, RCP6.0, and RCP8.5) revealed contrasting impacts of climate change and taxonomic reidentification on the future habitat availability of *R. dubia* and *R. indica* (Fig. 6c□h). For *R. dubia*, predicted niche shifts under future climate scenarios were relatively limited (Fig. 6c, e, and g). Only under the medium-emission scenario (RCP6.0) did new suitable areas emerge—particularly in southwestern China, spanning southern Sichuan, Yunnan, and Guizhou—which were apparent only after species reidentification (Fig. 6e), suggesting localized sensitivity to moderate climate change conditions. In contrast, *R. indica* exhibited substantial niche loss across all scenarios, particularly in central and southern China (Fig. 6d, f, and h). Notably, many of the lost niches were newly established following reidentification, emphasizing the ecological relevance of refined species boundaries and highlighting regions of emerging conservation concerns. These results demonstrate how both taxonomic accuracy and climate trajectories shape future species distributions and inform conservation prioritization.

## Discussion

### The 3D framework for delimiting taxonomically complex group

We introduced the 3D (Delimitation, Distribution, and Decoding) framework, a structured approach that synthesized data across spatiotemporal scales (Fig. 1). This involves fine-scale, population-level sampling for trait quantification (e.g., seed arrangement, petal number, genome size) under standardized conditions, as well as broad-scale, historical data derived from digital and physical herbarium specimens (Burbano & Gutaker 2023; Hong 2025). Taxonomic traits include not only classical diagnostic characters described in floras, but also anatomical and cytological features evaluated by their evolutionary relevance. Beyond controlled experiments, the framework also encourages trait assessments under seminatural conditions, such as common garden settings or field trials, to account for environmental effects and phenotypic plasticity.

By linking contemporary experimental evidence with historical collections and computational classification, the 3D framework supports robust secondary species delimitation, enhanced distributional accuracy, and feasible classification rationale underlying taxonomic assignments. The knowledge of morphological changes aligned with accurate species delimitation can further guide genetic studies of natural variation or wild relative-facilitated crop breeding, especially in polyploid systems (Dai et al. 2005; Castillo-Lorenzo et al. 2024; Han et al. 2024b; Xin et al. 2024). In addition, well-developed machine learning models from this framework may be used to retrain AI-based taxonomic tools, thereby improving their performance and consistency (Edwards & Knowles 2014; Karbstein et al. 2024). In summary, our integrative approach underscored the importance of accurate delimitation in polyploid complexes and provided foundational insights for subsequent ecological, evolutionary, and conservation studies.

### Integrative identification of morphologically overlapping polyploid complex

Accurate species delimitation within polyploid complexes is notoriously challenging due to extensive phenotypic plasticity and morphological convergence among taxa (Stebbins 1971; Hörandl 2022). Our integrative 3D approach, combining morphology, cytology, and phylogeny, significantly clarified species boundaries within the *Rorippa dubia–indica* complex (Fig. 2). Traits with high phylogenetic signals, particularly seed arrangement, petal number, and genome size, provided robust diagnostic markers that reflect underlying evolutionary histories (Tu et al. 2018; 2019). In contrast, leaf shape and fruit length demonstrated weaker phylogenetic signals, indicating these traits may represent environmentally driven plastic responses rather than stable taxonomic features (Chitwood & Sinha 2016).

Trait plasticity complicates field-based species delimitation (Hong 2025). The pronounced misidentification rates documented in initial field and virtual-herbarium assessments underscore the difficulty in distinguishing taxa using traditional morphological traits (Fig. 3) (Lukhtanov 2019; Wu et al. 2023). Original identifications often relied on traits with high plasticity (Rollins 1969; Gu & Hsu 1986), such as leaf morphology and fruit shape, which generated longer decision paths and higher ambiguity (Fig. 4). In contrast, anatomical traits like seed arrangement and petal number showing more evolutionarily conserved and discrete, enabled shorter, more precise classification pathways (Tu et al. 2019; Zheng et al. 2021). Minor inconsistencies were observed between phylogenetic placement and trait-based classification (occurrence rate = 1.2–1.8%; Fig. 2b), likely reflecting either incomplete lineage sorting, plastid capture, or developmental noises in certain polyploid individuals. This highlights the critical need for secondary assessments incorporating stable diagnostic traits to resolve initial taxonomic uncertainty (Wu et al. 2023; Chambers et al. 2025; Hong 2025).

Machine learning modeling reveals diagnostic complexity in taxonomy. The decision-tree approach provided valuable insights into the rationales underpinning taxonomic decisions (Smith & Carstens 2020; Sun 2025). Differences between original and second-round identification trees reflect shifts from reliance on plastic traits to more integrative trait-based criteria (Fig. 4). Deeper and more branched decision paths in the first round imply greater uncertainty and complexity in taxonomic reasoning due to the ambiguous nature of certain morphological traits. By contrast, the use of more conserved traits in the second round streamlined decision-making, enhancing classification efficiency and reproducibility. Hence, the complexity and depth of branching in decision trees offer a quantifiable measure of trait reliability and classification rationale in taxonomy (Edwards & Knowles 2014).

### Ecological implications of taxonomic refinement

Accurate delimitation also profoundly reshaped ecological interpretations and conservation priorities (Wu et al. 2023; Hong 2025). Kernel density analyses showed significant geographic shifts in species distributions following taxonomic revision (Fig. 5). Specifically, *R. dubia* became spatially cohesive, indicating previously fragmented occurrence data may have resulted from misidentifications. In contrast, *R. indica* displayed enhanced population isolation in central and southern China, highlighting previously unrecognized ecological differentiation. This improved spatial clarity has critical implications for conservation management, particularly under climate change scenarios (Hart et al. 2014; Hong 2016).

Species distribution models (SDMs) before and after reidentification underscored the ecological significance of accurate species boundaries. Following reassessment, predicted niches for *R. dubia* contracted notably around the middle-lower Yangtze River, whereas *R. indica* showed niche expansion into previously unrecognized suitable habitats across southeastern China (Fig. 6). Moreover, future climate scenario analyses revealed nuanced species-specific responses. *R. dubia* exhibited moderate vulnerability, with newly identified niches emerging only under the medium-emission scenario, highlighting localized sensitivity. Conversely, *R. indica* faced significant niche loss across central and southern China, particularly in newly established niche areas post-delimitation. Such differential climate sensitivities underscore the urgency of species-specific conservation strategies informed by refined taxonomy (Wu et al. 2023). Notably, many of the potentially lost habitats of *R. indica* overlapped with populations previously identified as having relatively small genome sizes (Han et al. 2024b). These accessions are of particular interest as genetic resources for *Brassica napus* improvement, given their demonstrated potential for distant hybridization across species boundaries (Dai et al. 2005; Xin et al. 2024). The projected loss of these populations underscores the urgency of targeted conservation efforts, especially for genetically valuable lineages that may be at heightened risk under future climate change.

### Broader implications for species conservation and research

This study emphasizes the profound consequences that taxonomic accuracy can have for ecological and conservation research within taxonomically complex polyploid species. Misidentifications can distort ecological modeling and conservation planning, potentially compromising the effectiveness of management interventions (Edwards & Knowles 2014; Hong 2016). The use of integrative 3D methods, combining stable morphological traits, phylogenetic insights, and cytological data, provides a robust framework to address these challenges (Wiens 2007; Karbstein et al. 2024). Future research should continue to refine and validate such approaches in other unresolved species complexes (Lukhtanov 2019; Hörandl 2022), improving taxonomic accuracy and ultimately supporting more effective conservation strategies.

## Author Contributions

T-SH conceived the project, collected the data, performed analyses and wrote the manuscript. J-XL did the specimen review and ecological analyses. Y-WX revised the manuscript. All authors reviewed the final manuscript.

## Acknowledgements

This work was supported by the National Natural Science Foundation of China (32170224 and 32225005) and the National Key Research and Development Program of China (2024YFF170140203). We acknowledge curators at the herbaria (IBSC, KUN, NAS, PE, and WUK) for assisting specimen review. We thank Quan-Jing Zheng and undergraduates for contributing to data collection.

## Data Availability

The sequences can be accessed by GenBank IDs: *Rorippa dubia* (PV802282-PV802362), *Rorippa hengduanshanensis* (PV802195-PV802208), and *R. indica* (PQ433226−PQ433289). Custom scripts and input data are available in the Dryad Digital Repository (http://datadryad.org/share/wDvUY6sW0zDP0XDJLBluogH_AuxM9_84WOfBqoBxZE4).

